## Supplementary figures for "Banp regulates DNA damage response and chromosome segregation during the cell cycle in zebrafish retina"

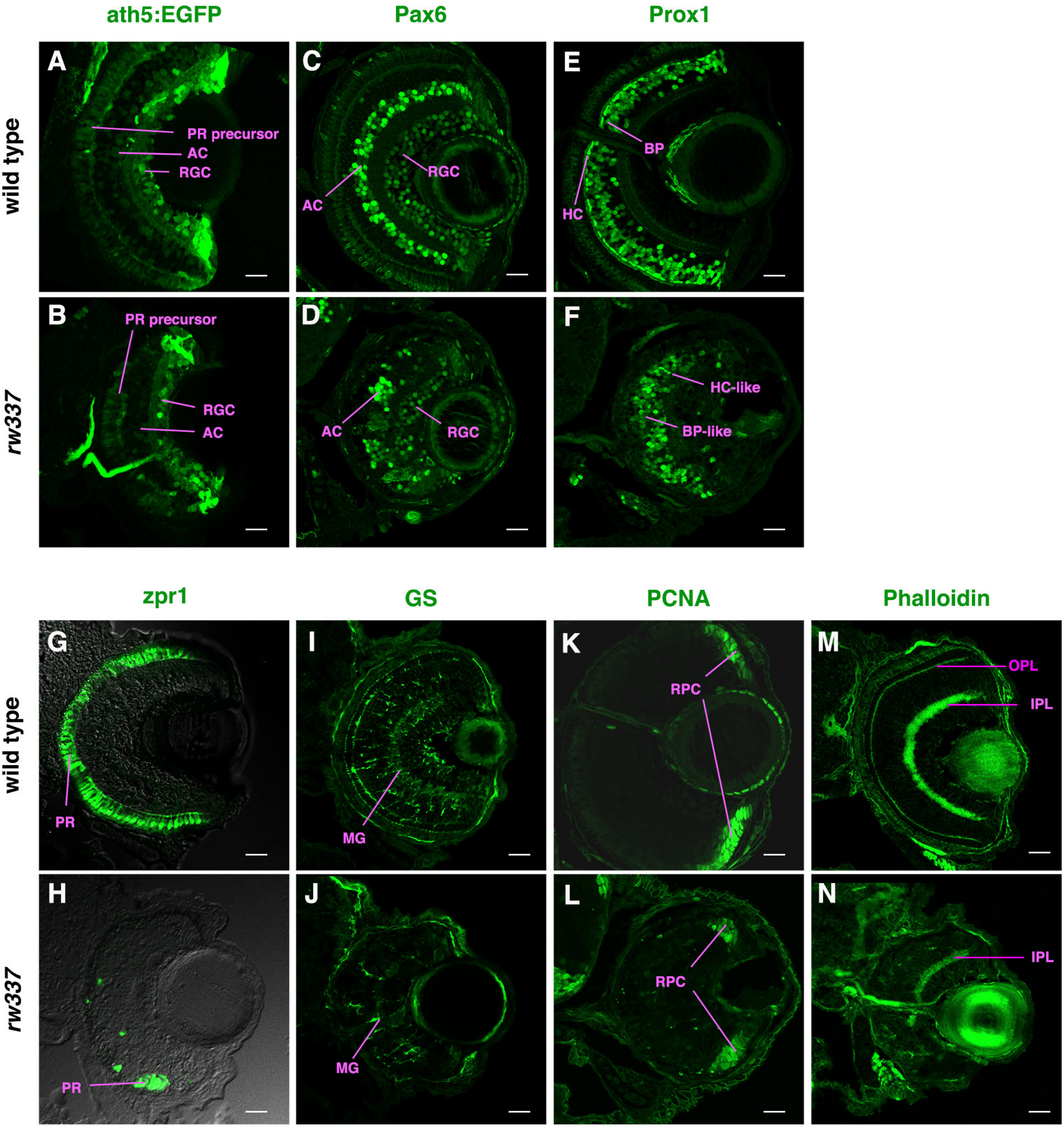


### Figure 1-figure supplement 1

#### Results colour-coded for amino acid conservation

The current colourscheme of the alignment is for **amino acid conservation.**

The conservation scoring is performed by PRALINE. The scoring scheme works from 0 for the least conserved alignment position, up to 10 for the most conserved alignm
The colour assignments are:

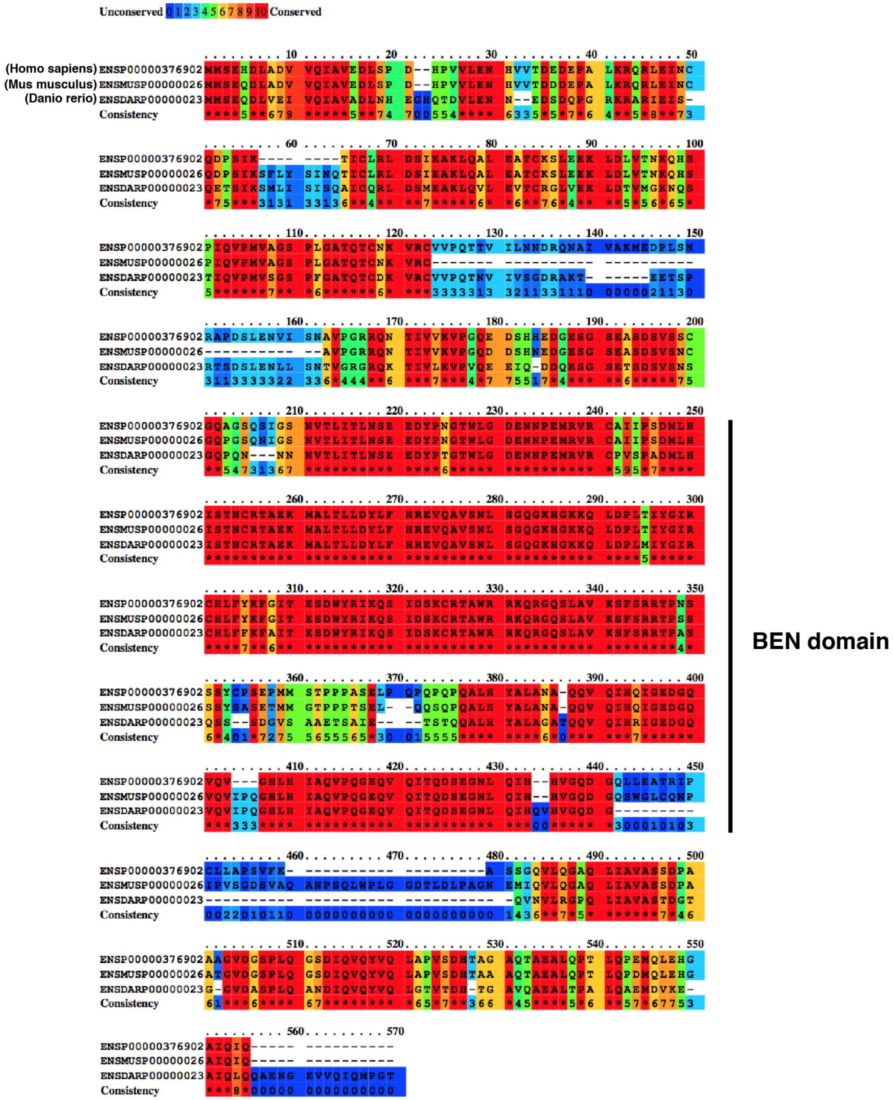


### Figure 1-figure supplement 2

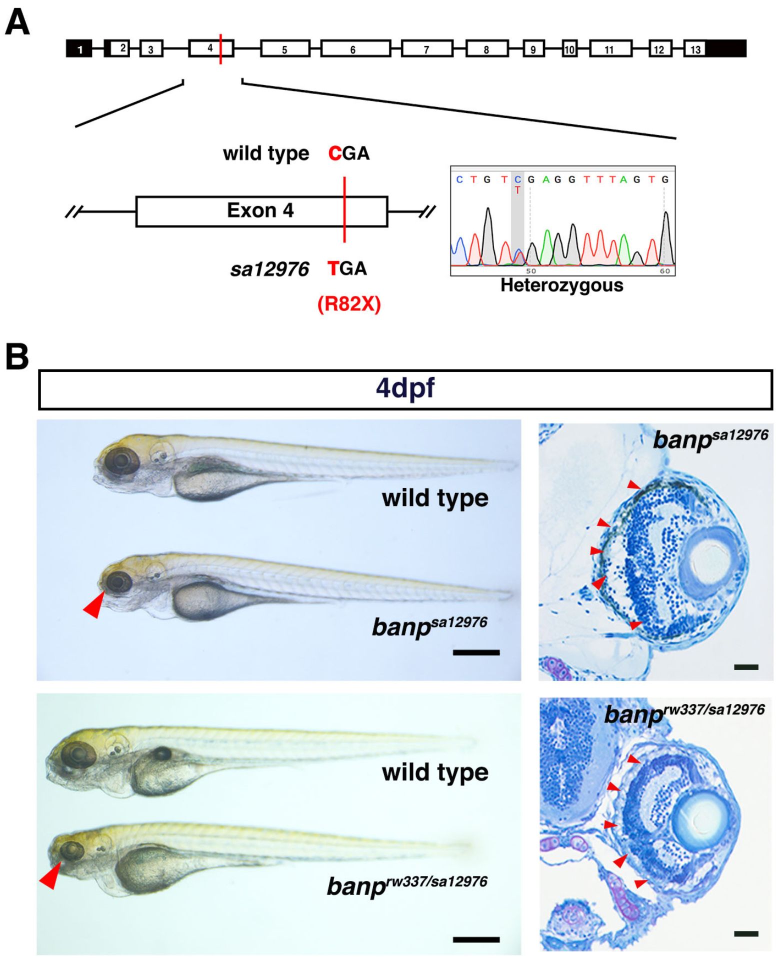


### Figure 1-figure supplement 3

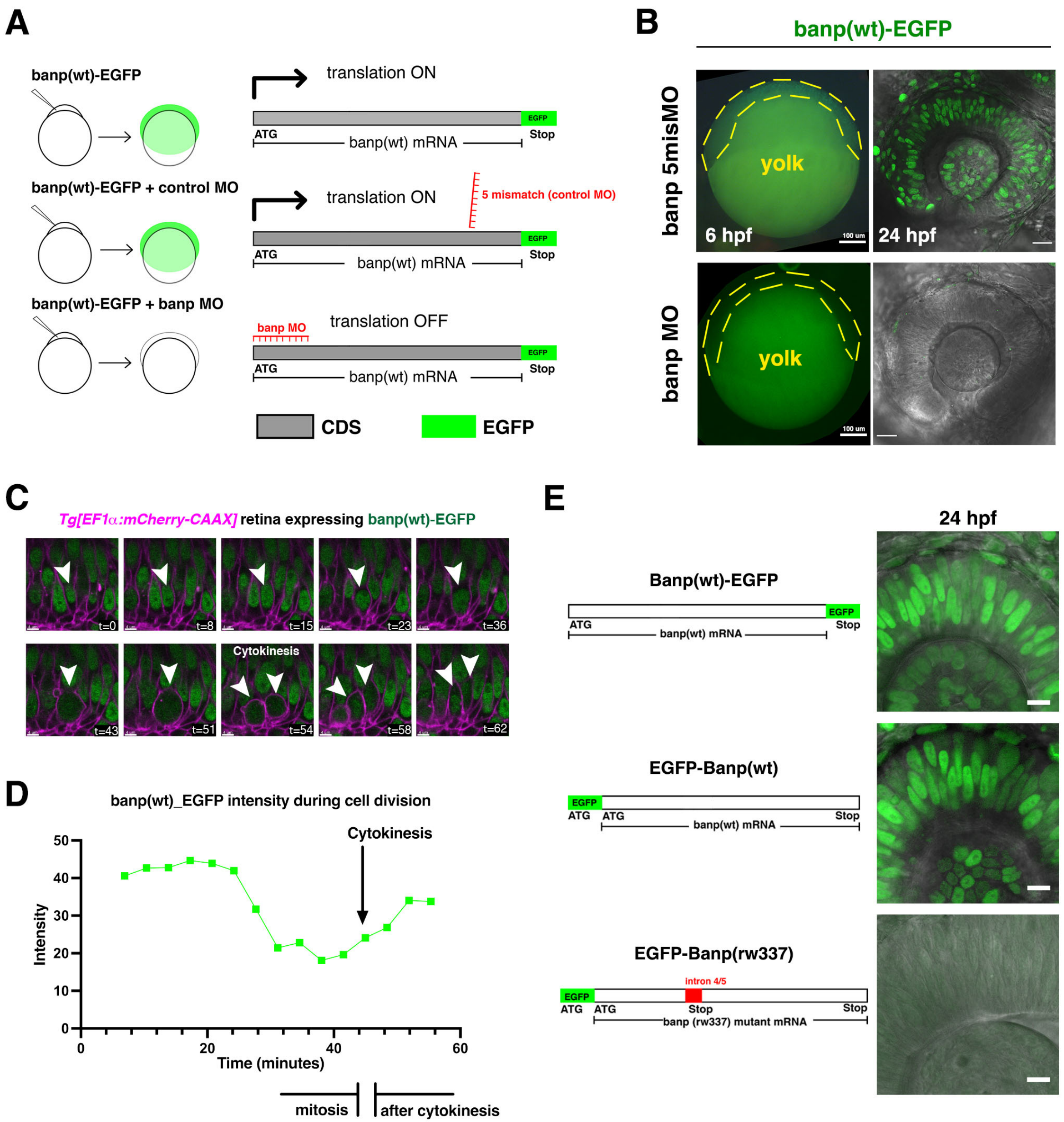


### Figure 1-figure supplement 4

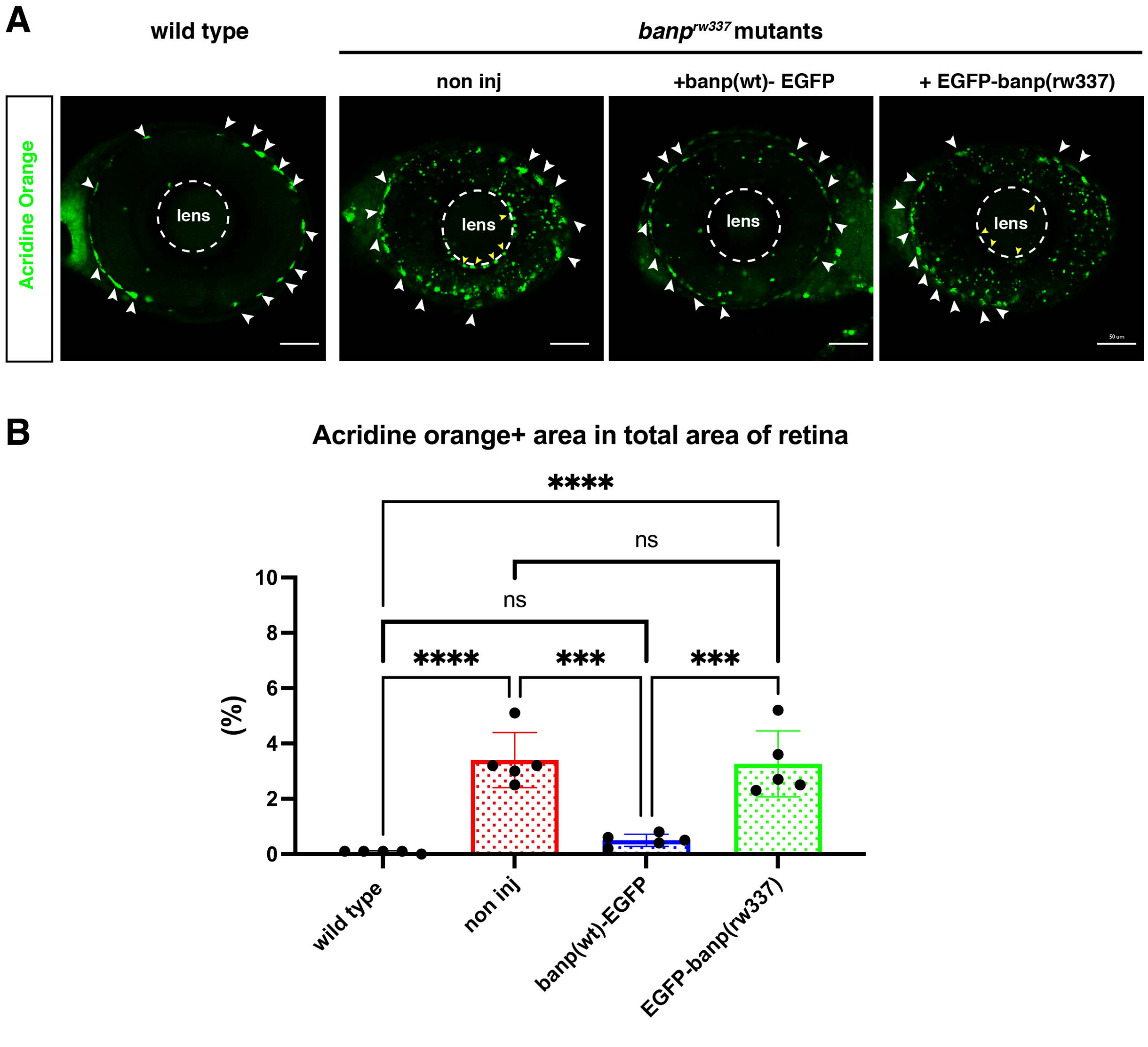


### Figure-1 figure supplement 5

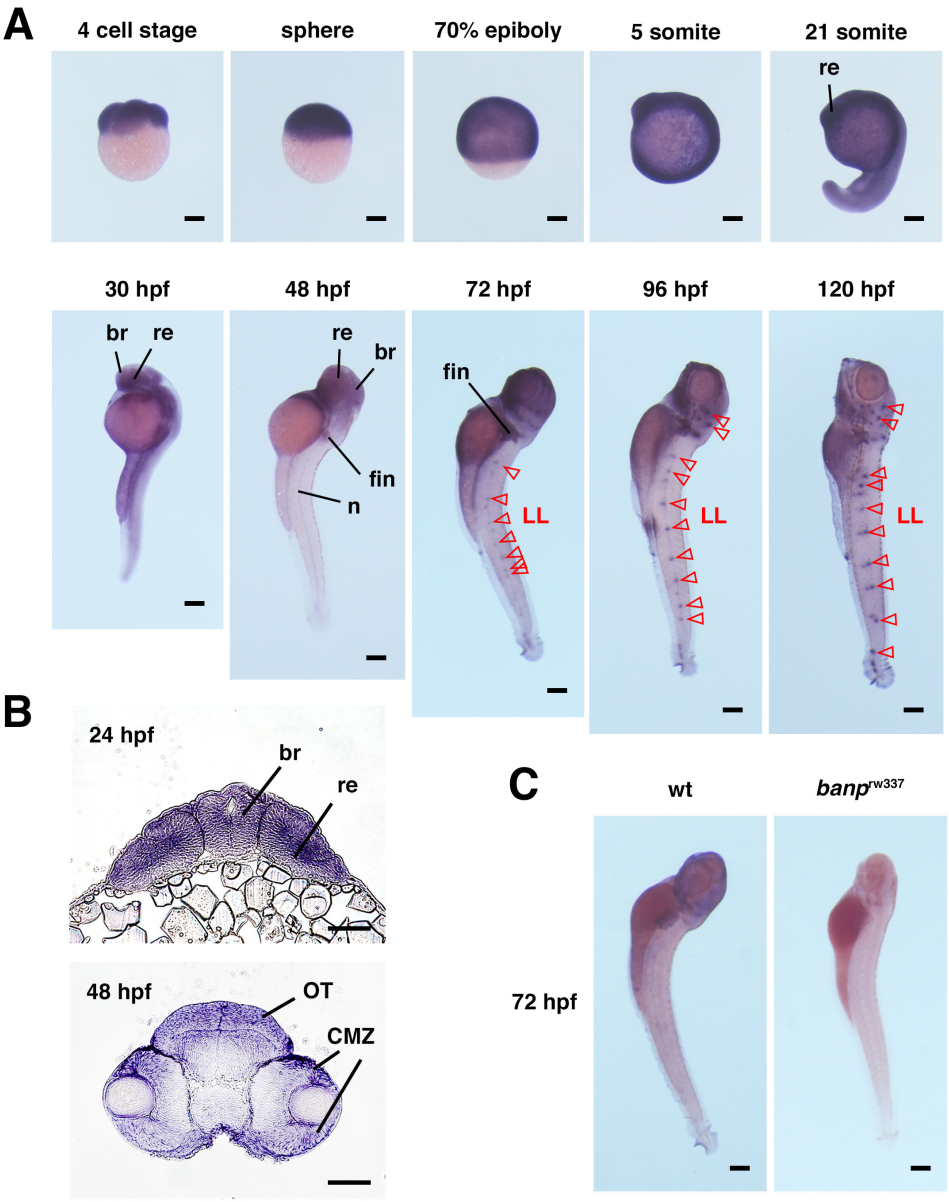


Fig. 1-figure supplement 6

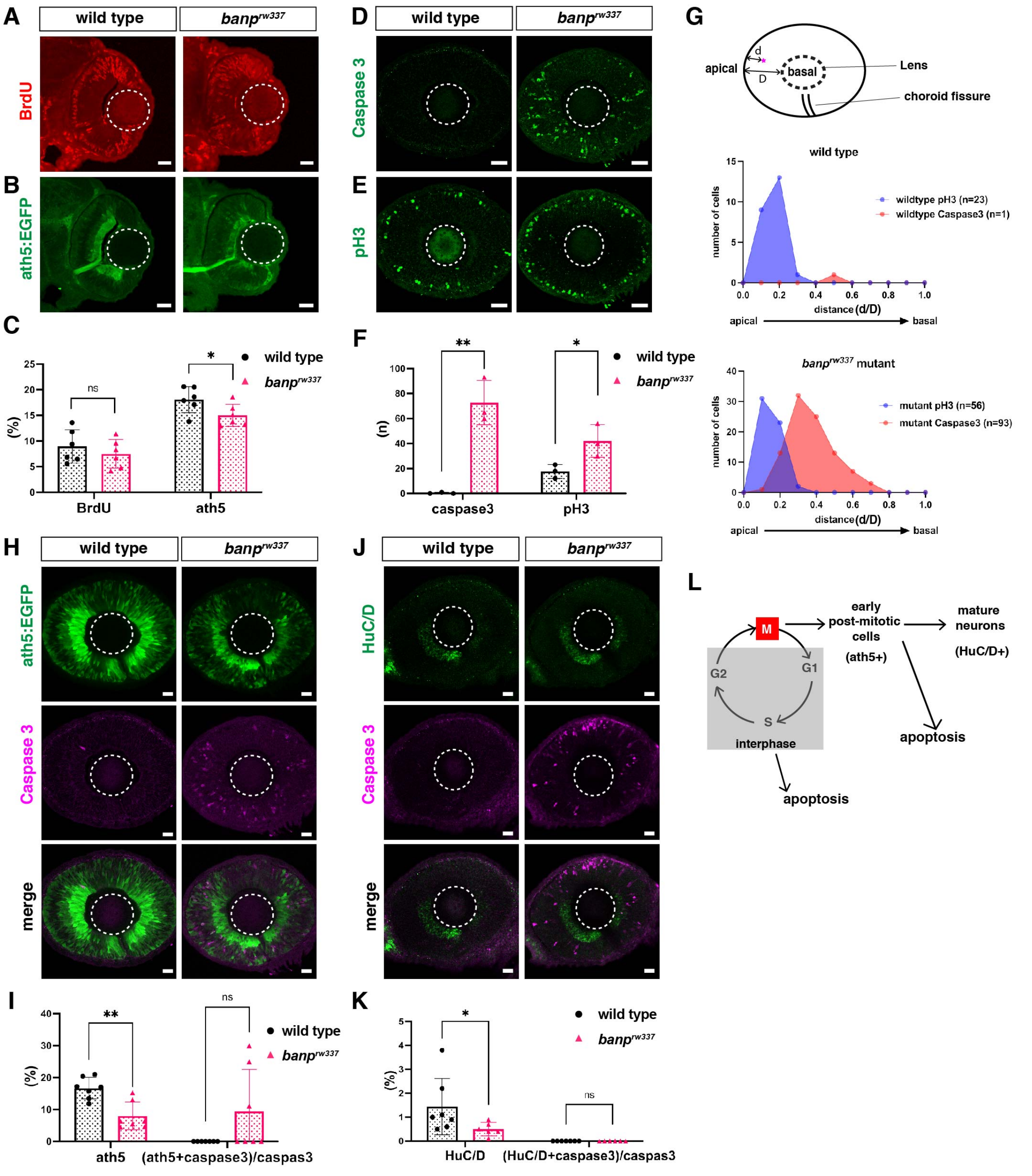


### Figure 2-figure supplement 1

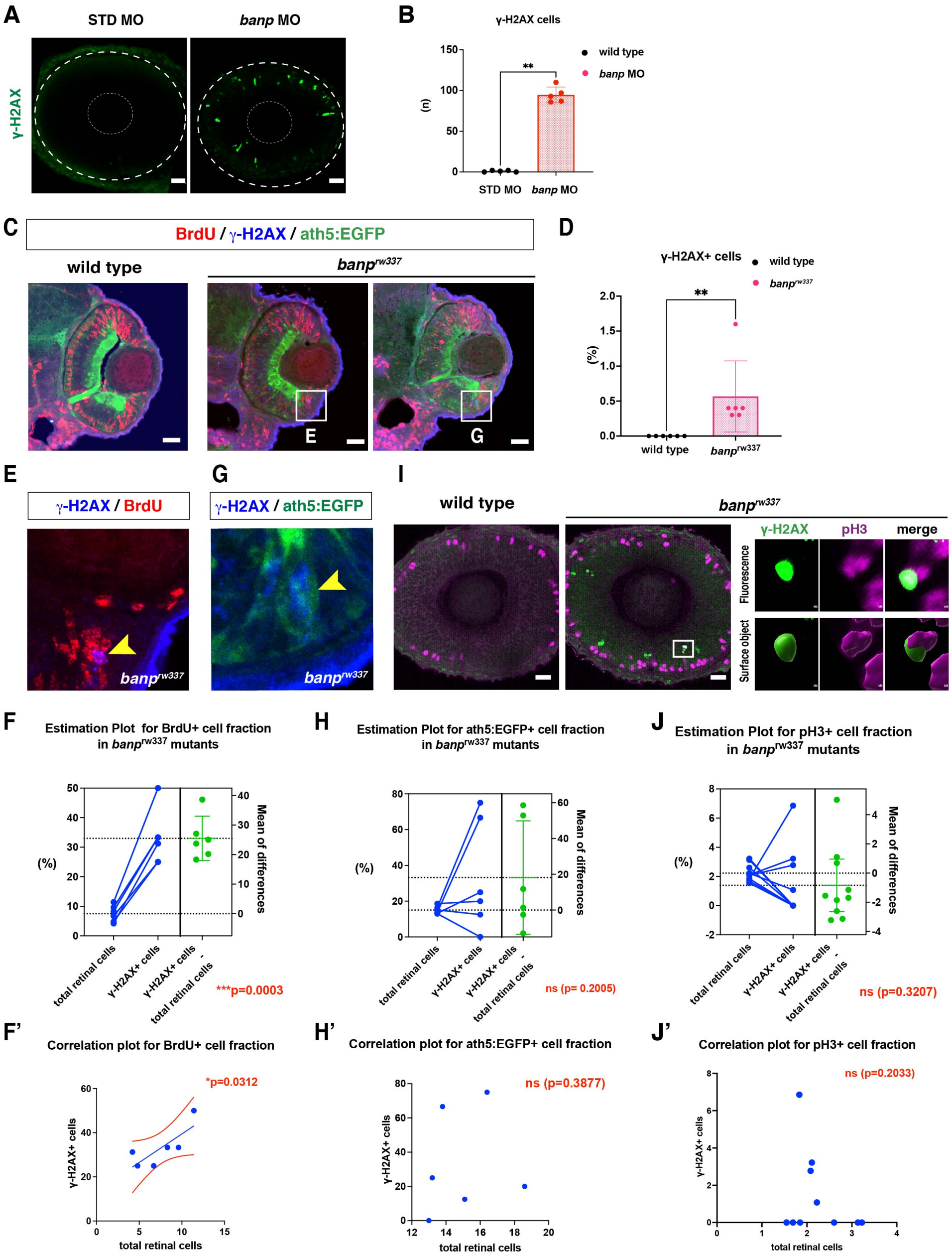


**Figure 4-figure supplement 1**

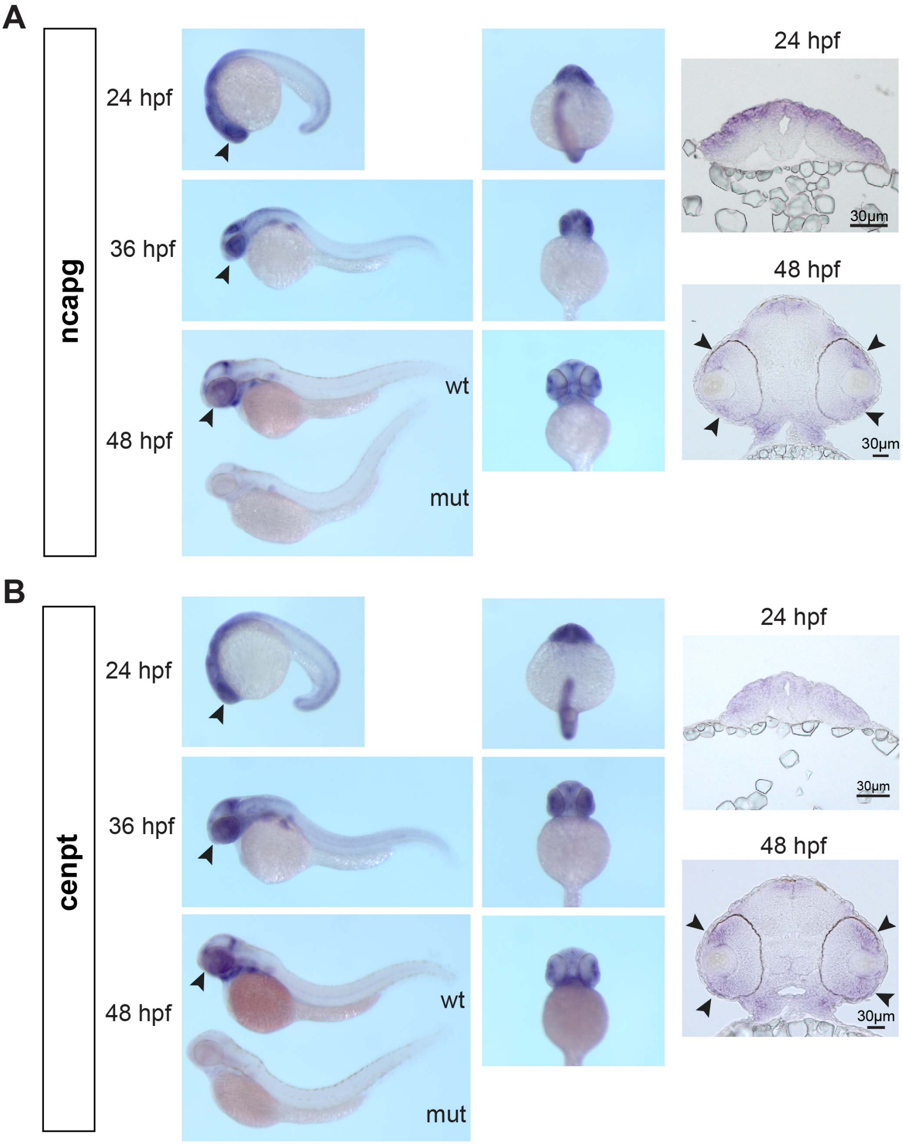


**Figure 5-figure supplement 1**

**A**

##### Top 6 downregulated motifs in banp$^{rw337}$ mutants

| Rank | Motif | P-value | log P-pvalue | % of Targets | % of Background | STD(Bg STD) | Best Match/Details |
|---|---|---|---|---|---|---|---|
| 1 | AGTCTCGCGAGA | 1e-58 | -1.339e+02 | 4.91% | 0.02% | 20.4bp (28.0bp) | GFX(?)/Promoter/Homer(0.941) More Information \| Similar Motifs Found |
| 2 | CTAATCCC | 1e-20 | -4.802e+01 | 9.66% | 2.12% | 25.6bp (30.9bp) | OTX2/MA0712.2/Jaspar(0.968) More Information \| Similar Motifs Found |
| 3 | CCCCTCAGCTGC | 1e-14 | -3.267e+01 | 0.98% | 0.00% | 17.8bp (0.0bp) | NHLH1/MA0048.2/Jaspar(0.706) More Information \| Similar Motifs Found |
| 4 | ACAACCAATCAG | 1e-13 | -3.206e+01 | 4.58% | 0.69% | 23.3bp (25.4bp) | NFYA/MA0060.3/Jaspar(0.936) More Information \| Similar Motifs Found |
| 5 | CGGCGCCATCTT | 1e-12 | -2.913e+01 | 5.89% | 1.32% | 29.6bp (27.0bp) | YY1(Zf)/Promoter/Homer(0.857) More Information \| Similar Motifs Found |
| 6 | TGGAGTGGCAGC | 1e-12 | -2.888e+01 | 1.64% | 0.04% | 24.9bp (25.8bp) | Pbx3(Homeobox)/GM12878-PBX3-ChIP-Seq(GSE32465)/Homer(0.795) More Information \| Similar Motifs Found |

##### Top 6 upregulated motifs in banp$^{rw337}$ mutants

| Rank | Motif | P-value | log P-pvalue | % of Targets | % of Background | STD(Bg STD) | Best Match/Details |
|---|---|---|---|---|---|---|---|
| 1 | TCATTAATCACT | 1e-23 | -5.522e+01 | 8.37% | 1.03% | 28.1bp (30.1bp) | Pdx1(Homeobox)/Islet-Pdx1-ChIP-Seq(SRA008281)/Homer(0.917) More Information \| Similar Motifs Found |
| 2 | GAGTGTTGGGGA | 1e-15 | -3.497e+01 | 4.98% | 0.57% | 23.6bp (26.5bp) | HNRNPK(KH)/Homo_sapiens-RNCMPT00026-PBM/HughesRNA(0.687) More Information \| Similar Motifs Found |
| 3 | GGGGGAGCCTCT | 1e-14 | -3.399e+01 | 1.20% | 0.00% | 29.4bp (14.1bp) | GCR1/Literature(Harbison)/Yeast(0.696) More Information \| Similar Motifs Found |
| 4 | AGCAAGCGAC | 1e-12 | -2.960e+01 | 9.16% | 2.53% | 27.7bp (28.9bp) | MBNL1(Znf)/Homo_sapiens-RNCMPT00038-PBM/HughesRNA(0.841) More Information \| Similar Motifs Found |
| 5 | TCCATTCGGT | 1e-12 | -2.914e+01 | 4.78% | 0.68% | 25.4bp (27.2bp) | TARDBP(RRM)/Homo_sapiens-RNCMPT00076-PBM/HughesRNA(0.661) More Information \| Similar Motifs Found |
| 6 | CAGCCTGTCACA | 1e-12 | -2.801e+01 | 2.99% | 0.22% | 25.5bp (30.7bp) | Hth/dmmpmm(Noyes_hd)/fly(0.649) More Information \| Similar Motifs Found |

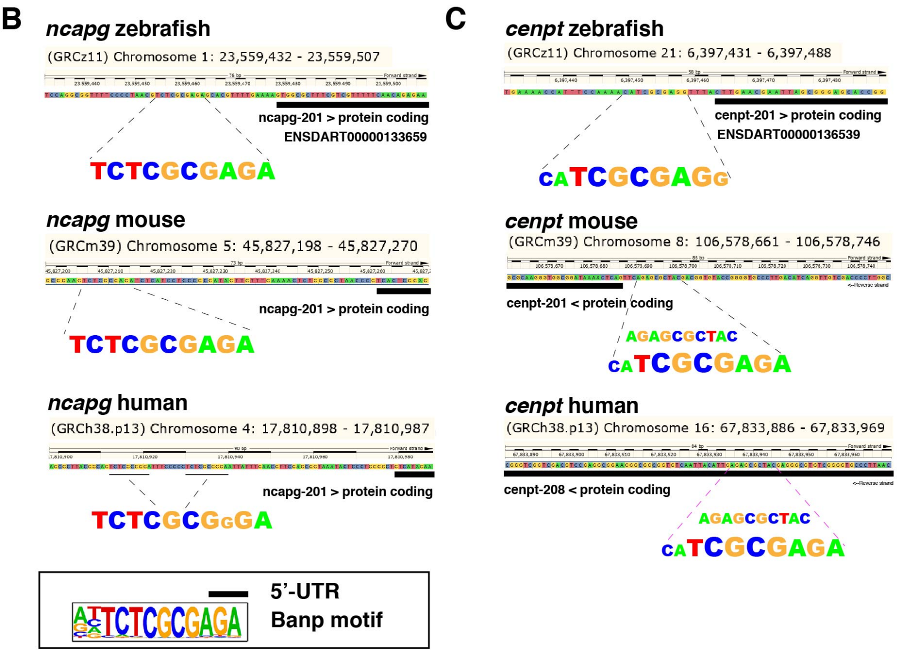


#### Figure 6-figure supplement 1

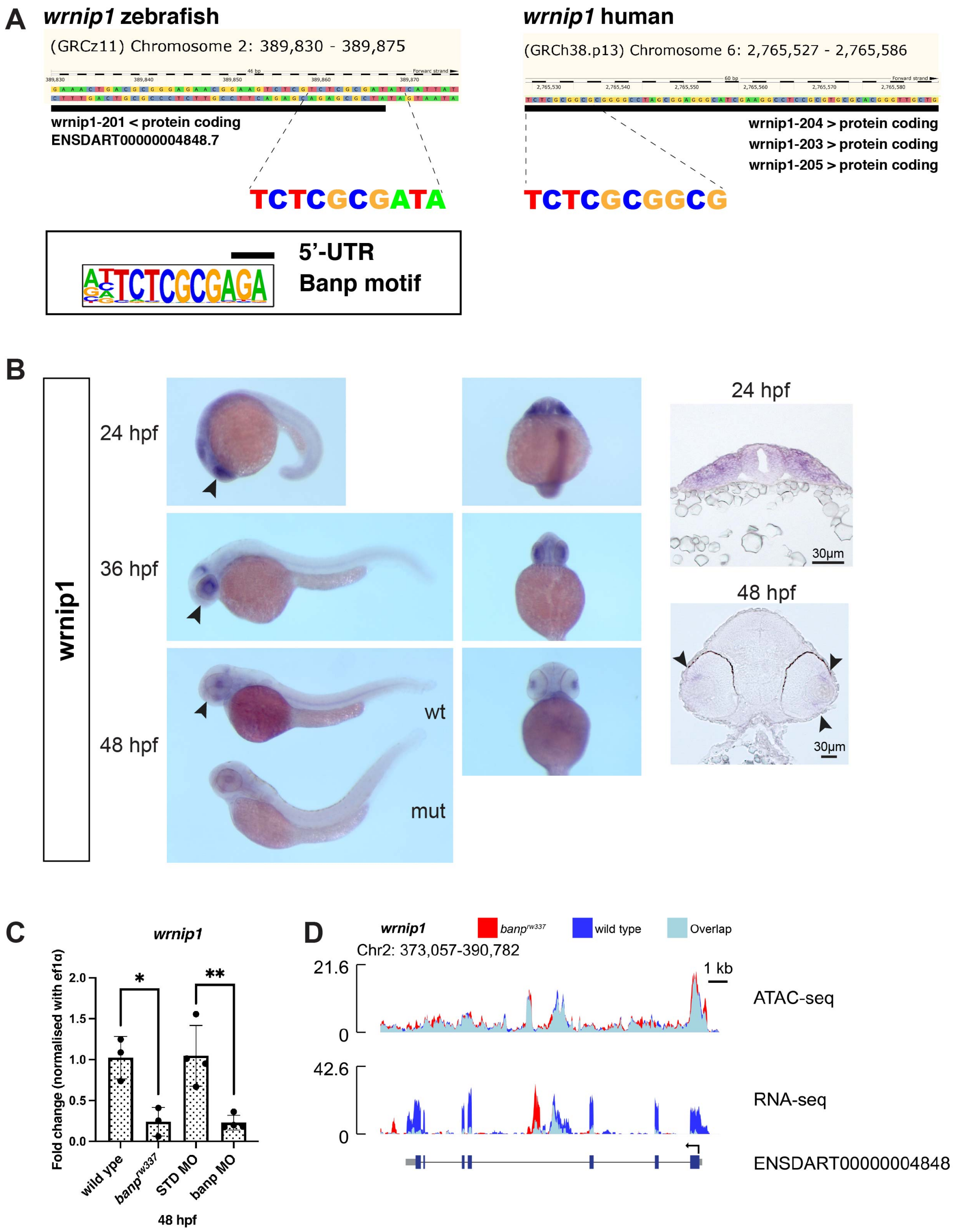


**Figure 6-figure supplement 2**
